# Estimating Effects Of Second Line Therapy For Type 2 Diabetes Mellitus: Retrospective Cohort Study

**DOI:** 10.1101/130724

**Authors:** Assaf Gottlieb, Chen Yanover, Amos Cahan, Yaara Goldschmidt

## Abstract

**Objective:** Metformin is the recommended initial drug treatment in type 2 Diabetes Mellitus, but there is no clearly preferred choice for an additional drug when indicated. We use electronic health records to infer the counterfactual drug effectiveness in reducing HbA1c levels and effect on body-mass index (BMI) of four second line diabetes drug classes.

**Study design and setting:** Retrospective analysis of the electronic health records of US-based patients in the Explorys database using causal inference methodology to adjust for censored patients and confounders.

**Participants and Exposures:** Our cohort consisted of roughly 25,000 patients with type 2 diabetes, prescribed metformin along with a drug out of four second line drug classes – sulfonylureas, thiazolidinediones, DPP-4 inhibitors and GLP-1 agonists, during the years 2000-2013.

**Main outcome measures:** Glycated hemoglobin (HbA1c) and BMI of these patients after six and twelve months of treatment.

**Results:** We show that all four drug classes reduce glycated hemoglobin levels, but the effect of sulfonylureas after 12 months of treatment is less pronounced compared to other classes. We also predict that thiazolidinediones increase body weight while DPP-4 inhibitors decrease it.

**Conclusion:** Our results are in line with current knowledge on second line drug effectiveness and effect on BMI. They demonstrate that causal inference from Electronic health records is an effective way for conducting multi-treatment causal inference studies.

### WHAT IS ALREADY KNOWN ON THIS TOPIC

The effect of type 2 diabetes second-line drugs on glycosylated hemoglobin levels and on Body-Mass Index have been evaluated in clinical studies. However, the clinical implication of these studies is limited by small number of participating individuals and the homogeneity of the study populations. Meta-analysis studies have increased sample size but potentially suffer from the similar homogeneity biases.

### WHAT THIS STUDY ADDS

This study performs, for the first time, a large-scale analysis of the therapeutic and adverse effects of type 2 diabetes second-line drugs in real-world population using electronic health records. We confirm current knowledge for glycosylated hemoglobin levels, while showing similar effects on BMI for inhibitors of Dipeptidyl peptidase 4 (DPP-4), Glucagon-like peptide-1 receptor agonists (GLP-1).

## Introduction

Type 2 Diabetes Mellitus (T2DM) affects more than 29 million people in the United States and is the 7^th^ leading cause of death (1),(2). The American Diabetes association standards of medical care (3), supported by several studies (4,5), recommends dietary changes and physical exercise as the initial treatment, followed by administration of metformin if life style changes fail to reach glycemic control. According to the standards of medical care, if metformin does not achieve glycemic target within three months, one of the following six second-line medications should be added: Sulfonylureas (SU), thiazolidinediones (TZD), inhibitors of Dipeptidyl peptidase 4 (DPP-4), Glucagon-like peptide-1 receptor agonists (GLP-1), SGLT2 inhibitors, or insulin. Currently, the guidelines do not prefer one class over the others. The effectiveness, costs and risk of complication of those drug classes were compared in clinical trials (6) and meta-analyses of their results (7–9). These comparisons found no significant difference in drug class effect on the percentage of blood glycated hemoglobin (HbA1c), thus no specific recommendation about the choice of a second drug could be made (10).

Notably, clinical trials are laborious and costly. Trials often include small samples with limited representativeness of the target population (e.g., between 2005 and 2012, the FDA approved drugs based on a median number of two clinical trials and median number of patients enrolled was 760 (11)). Meta-analyses of clinical trials may have higher power and be more generalizable, but are vulnerable to publication bias, small-study effects and limited degree of heterogeneity (12). Electronic health records (EHRs) hold promise as an alternative way to conduct causal inference experiments, that can address some of these meta-analyses limitations (13,14).

Here, we demonstrate the usefulness of a real world evidence approach for T2DM. We simulate a multi-arm clinical trial of four classes of drugs for diabetes which are commonly used as second line treatment (SU, TZD, DPP-4 and GLP-1) and compare the counterfactual effectiveness (HbA1c levels) and body-mass index (BMI) outcomes of 25,098 patients over the course of twelve months, adjusting for confounders and censored patients. For reference, a recent meta-analysis of anti-diabetes drugs (8) was based on data of about 18,000 patients. Our results are in line with current knowledge and thus demonstrate that causal inference from EHRs is an effective way for conducting multi-treatment causal inference studies.

## Research Design and Methods

### Data

We used the Explorys database (IBM Inc.), which includes EHR records of 47 million patients, pooled from multiple different healthcare systems in the US. Data consists of a combination of clinical EHRs, healthcare system outgoing bills, and adjudicated payor claims and is standardized and normalized using common ontologies, searchable through a HIPAA-enabled, de-identified database tools. The EHR data includes patient demographics, diagnoses, procedures, prescribed drugs, vitals and laboratory values.

### Cohort definition

We defined a cohort of T2DM patients based on the Northwestern University diabetes phenotyping algorithm (15). We included patients having at least two types of evidence for T2DM out of a T2DM diagnosis, T2DM drugs and indicative lab values (glucose or HbA1c levels). Included were patients who were first prescribed metformin and subsequently prescribed, during the years 2000-2013, a second line drug belonging to any of four classes: sulfonylureas (SU), thiazolidinediones (TZD), inhibitors of dipeptidyl peptidase 4 (DPP-4) and Glucagon-like peptide-1 receptor agonists (GLP-1) (Table S1 lists drugs for each drug class). The first completed prescription of the second-line drug was considered the “index-date”. Drug combinations of two or more second line drug classes were considered as prescription of the two drug classes at the same day. Patients prescribed two or more second line drugs were censored (see “causal inference scheme” for further details).

We required the patients to have at least twelve months of documented observation period prior to the index-date and a documented fifteen months of observation for the follow-up period. Additionally, we required each patient to have at least one HbA1c measurement during the observation period and at least one measurement for each of the follow-up time points of six and twelve months (±three months each) from the index-date. Finally, we excluded patients with type 1 Diabetes Mellitus, identified by either a type 1 Diabetes diagnosis code, or prescription of pramlintide (approved also for T2DM patients who use insulin).

### Feature extraction

We extracted patient characteristics using the feature engineering framework of (16). The comprehensive set of features included demographic information (age, sex, ethnicity), insurance type, patient aggregated diagnoses using Clinical Classifications Software (CCS) categories, Charlson comorbidity index (CCI) (17) and Elixhauser comorbidity index categories (18), prescribed drugs (active ingredients) and laboratory results values over the baseline period. Categorical features, such as insurance type or ethnicity, were split into binary features. As a preliminary step, we filtered features which were dominated (>95% of patients) by a single value or were spurious (>80% with missing values), resulting in 523 features.

### Causal inference scheme

We considered two potential biases: (1) selection bias due to censoring; and (2) confounders, affecting both treatment choice and measured outcome (HbA1c levels or BMI).

For HbA1c inference, we marked as censored patients who, during the follow-up period, were switched to or added any other anti-diabetic drug (including classes which were not evaluated in this work: insulin, sodium-glucose co-transporter 2 inhibitors, meglitinides and α-glucosidase inhibitors). We also censored patients who underwent bariatric surgery during the follow-up period. For BMI prediction, we followed the same censoring criteria as in HbA1c inference and additionally censored patients lacking BMI measurements in either of the two follow-up time points of six and twelve months. We corrected for censoring by re-weighing the uncensored population using inverse probability weighting (IPW) (19).

We defined the set of confounders in two ways: (1) confounders identified by an internist and through literature search; and (2) treating all 523 extracted features as confounders. In total, 30 domain-expert confounders for HbA1c inference (Table S2) and 6 domain-expert confounders for BMI inference (Table S3) were identified. We corrected for confounders using either standardization or stabilized inverse probability weighting (IPW) (20). For standardization, we used ridge regression with five-fold cross validation to adjust the regularization coefficient and for the inverse probability weighting, we used multi-class logistic regression.

We checked the balancing of confounders in our IPW scheme by applying the diagnostics of (21). We further applied our scheme to patient height as a negative control, a measurement that is unaffected by treatment type.

### Patient involvement

No patients were involved in setting the research question or the outcome measures, nor were they involved in developing plans for design or implementation of the study. No patients were asked to advise on interpretation or writing up of results. There are no plans to disseminate the results of the research to study participants or the relevant patient community.

## Results

### Study cohort

Our cohort included 25,098 patients. Of these, 16,327 also had available BMI before and after the prescription of second line drugs and were used for inference of BMI (treating the rest as censored) (Table 1). There were significantly more patients on TZD, GLP-1 or DPP-4 that switched or added another drug than patients on SU (censored patients, p<9e^-4^, Table 1). TZD and SU had significantly higher percentage of patients with missing BMI measurements during the follow-up than GLP-1 and DPP-4 (p<8e^-6^, Table 1). Finally, the patients on GLP-1 were about five years younger on average (p<4e^-57^) and included significantly higher rate of women (p<2e^-24^, Table 1).

**Table 1.**
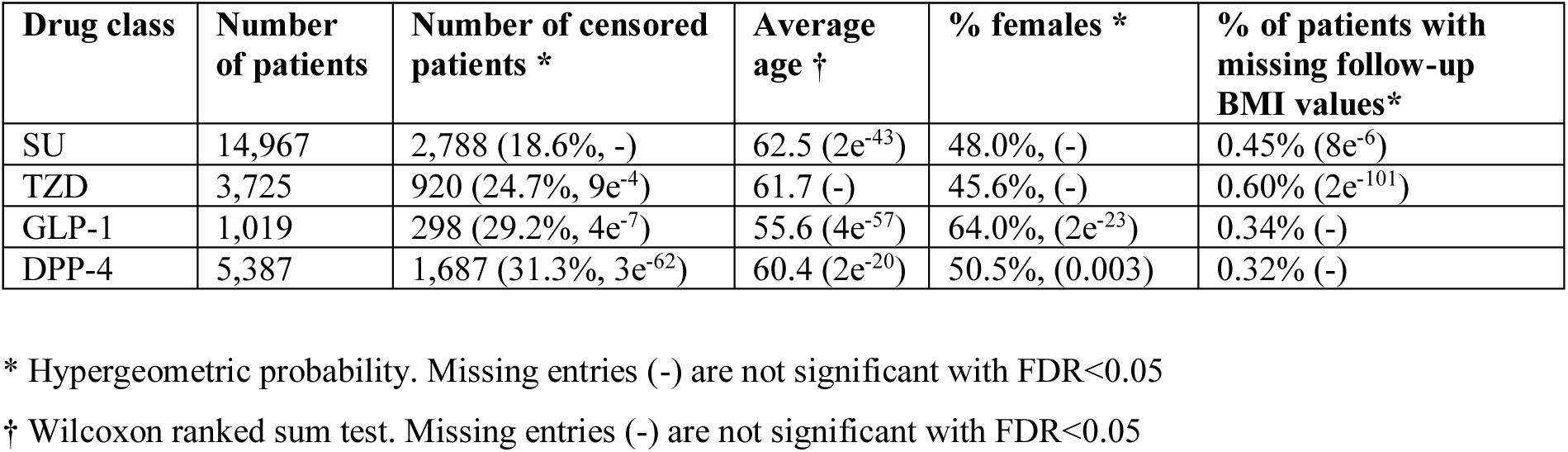
Descriptive statistics of patients on T2DM second line drug classes.

### Causal inference scheme

We applied causal inference methods to compute the counterfactual HbA1c levels and BMI (for each one of the four drug classes) at each of the two follow-up time points, adjusting for censored patients and confounders (Methods). Using height as a negative control, we found no significant differences between patients on different drug classes.

### HbA1c

HbA1c measurements were available for 82% of the patients from up to 90 days prior to initiation of second-line treatment, and for 94% of the patients up to 180 days (see Figure S1 for complete temporal distribution). The counterfactual HbA1c levels were significantly correlated with actual HbA1c levels in both time points (Pearson ρ≥0.32, p<e^-36^ using domain expert confounders and ρ≥0.41, p<e^-50^ using the comprehensive confounder set, Table S4).

The differences in predicted HbA1c levels between causal inference methods (standardization and IPW (19)) and between manual and automatic selection of confounders were lower than 0.2% and significant only between causal inference methods for SU patients (Wald test, below false discovery rate, FDR, of 0.05, Table S5). All four drug classes were predicted to reduce HbA1c levels below 7.08% after twelve months of treatment, with a 0.18% (GLP-1) to 0.64% (SU) reduction in HbA1c levels relative to baseline (Table S5, Figures 1, S2). Twelve months HbA1c levels inferred for SU were higher than for TZDs, DPP-4 and GLP-1 by 0.05%-0.21%, significant under standardization reweighting (Wald test, p<3e^-5^, Table S6). Notably, both actual and inferred HbA1c levels were lower than those computed using the mixed-treatment comparison (MTC) of clinical trials of McIntosh et al (7,8).

**Figure 1.**
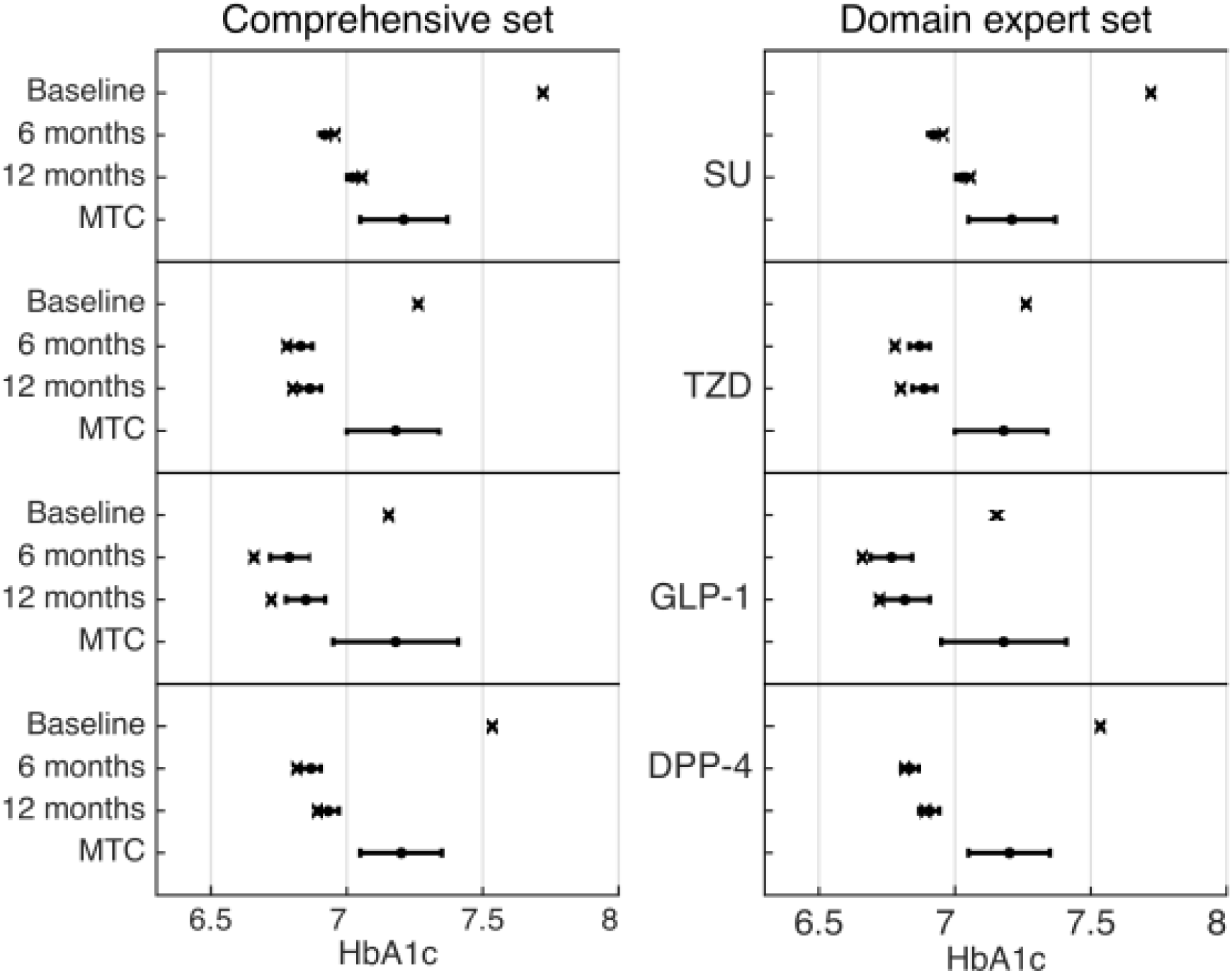
Predicted and observed HbA1c levels using standardization adjusting for either a comprehensive set of confounders (Left panel) or a set of confounders provided by a domain expert (Right panel). X marks indicate the actual measurements of patients at baseline (before second-line treatment), after six and twelve months. Black dots (with error bars) represent the counterfactual predictions and 95% confidence intervals, supposing all patients were treated with that drug class. The results of the Bayesian mixed-treatment comparison meta-analysis by McIntosh et. al (7,8) are marked MTC.

### BMI

BMI measurements were available for 77% and 82% of the patients as recent as 90 and 180 days prior to treatment date, respectively (see Figure S3 for complete time distribution). Counterfactual BMI was highly correlated with the actual BMI in both time points (ρ≥0.78, p<e^-97^).

The predicted BMI was significantly lower for patients on DPP-4 than for patients on SUs or TZDs (p<0.02, Figures 2, S4; Tables S7-S8). On average, patients on GLP-1 had higher BMI (by 2.5-2.7 kg/m^2^, Table S7) before the prescription of second line treatment compared to patients prescribed one of the other studied drug classes. These patients had their BMI lowered by an average of 0.74 kg/m^2^. However, our predicted BMI shows no significant advantage for GLP-1 over TZD and SU in a population with lower initial BMI (Table S8, see also discussion on BMI and GLP-1).

**Figure 2.**
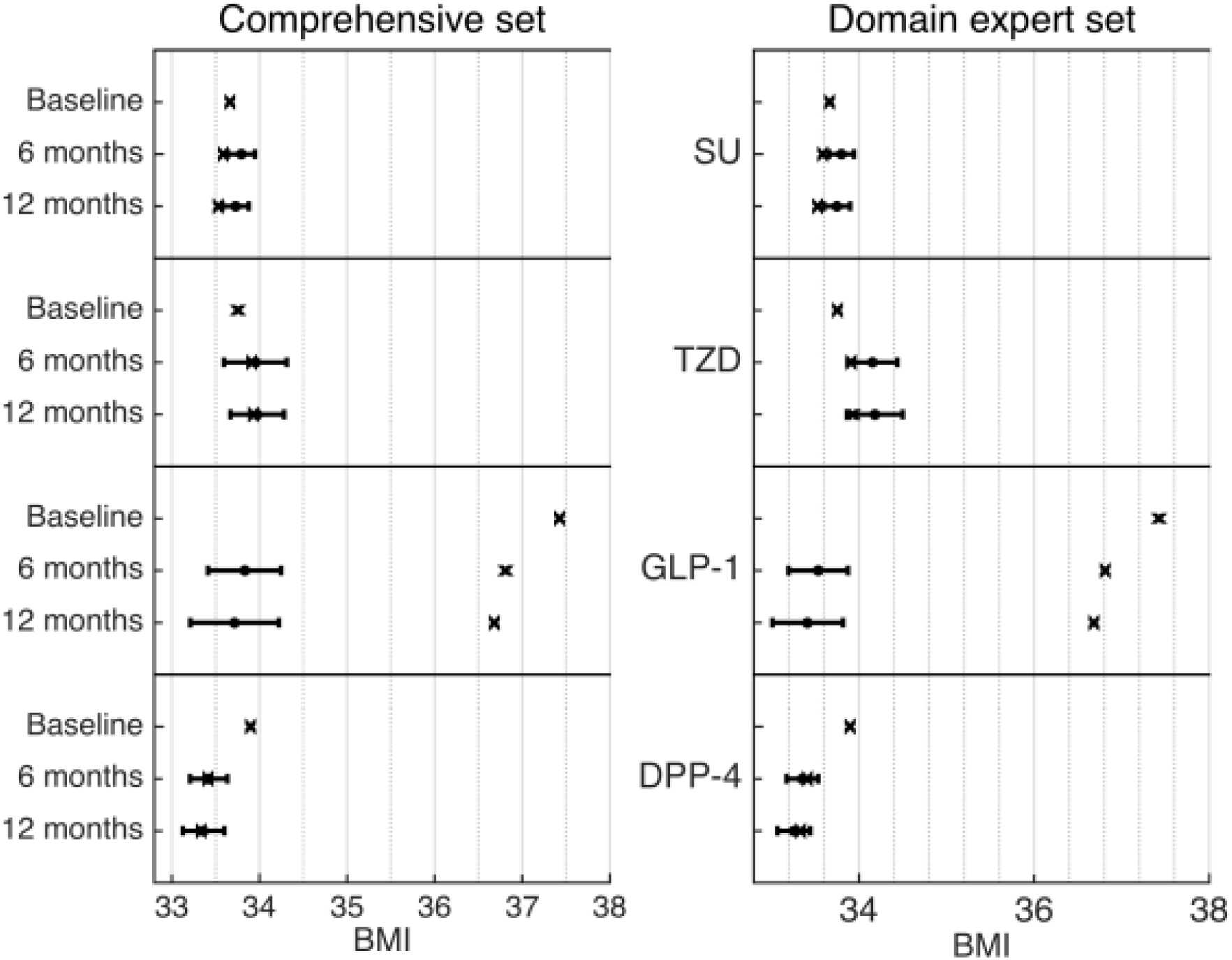
Predicted and observed BMI levels using standardization adjusting for either a comprehensive set of confounders (Left panel) or a set of confounders provided by a domain expert (Right panel). X marks indicate the actual measurements of patients at baseline (before second-line treatment), after 6 and 12 months. Black dots (with error bars) represent the counterfactual predictions and 95% confidence intervals, supposing all patients were treated with that drug class.

## Discussion

We presented a causal inference analysis of EHR data to compare the effect of adding a second line treatments for T2DM on HbA1c and BMI, in patients already treated with metformin. Our inferred HbA1c levels for up to twelve months of follow-up suggest that the effect of TZD, DPP-4 and GLP-1 inhibitors is comparable, whereas that of SU is smaller. At the same time, TZD increase BMI, whereas DPP-4 and GLP-1 reduce BMI and SU have a negligible effect on it after 12 months of treatment.

Our study has several limitations. We note three potential challenges encountered when performing causal inference with EHR data: (1) missing information for patients seen in clinics which are not included in the Explorys database; (2) incomplete tracking of prescriptions, leading to incomplete information about treatment changes; and (3) potential confounders not available in EHR data, such as life style. We consider HbA1c and BMI as good proxies for future patient risk (22), but there are other considerations in selecting a second line drug beyond its effect on these measures, such as risks of adverse reactions and of diabetes-related complications. While we did not directly address adverse reactions, patients who were switched drug classes may indirectly point to such effects. These outcomes should be studied in subsequent work, potentially observing patients for longer follow-up periods to gain stronger statistical power. Other extensions should focus on differences between individual drugs from the same class, which could have different outcomes (such as different drugs from the SU class (23)).

Patients prescribed SU were less likely to be added a third drug or switched to another drug than patients on the other drugs studied. This, despite the effect of SU on HbA1c being somewhat smaller. Possible explanations for this observation may include the low cost of SU, the availability of a metformin-SU pill (24) and the option of once-a-day dosing. Additionally, SU and TZD patients had significantly lower availability of BMI measurements during the follow-up period. Possible explanations for this are that when GLP-1 or DPP-4 treatments are prescribed, either the physician or the patient is more likely to have been concerned with the BMI, thus measures it more frequently; or that costs of these drugs tend to be lower and would be more frequently prescribed to patients with a lower socioeconomic state, which tend to be less well followed up on.

GLP-1 agonists are prescribed significantly more to women. Difference between men and women response to GLP-1 was reported in 2005 (25) and a study from 2013 found that the effect of one such GLP-1 agonist, exanatide, was larger in women (26). All patient in our study were treated with GLP-1 after 2005 and 58% of them treated after 2013, suggesting physicians may have considered this evidence when prescribing GLP-1. Also, patients on GLP-1 are typically younger than patients on other drug classes, in line with an observation made by others (27). Finally, patients with higher BMI tend to be prescribed GLP-1, and this is likely due to its known positive effect on weight (28,29).

TZD is the only class to maintain HbA1c at a stable level in six and twelve months, whereas HbA1c levels are increased over time in the other studied classes. A gradual weaning of the effect of SU on HbA1c levels had been previously described (13).

While we found good correspondence between standardization and IPW estimates in most of our analyses, the two differ significantly on BMI predictions for GLP-1. We suspect that the IPW corrections in this case were more susceptible to small sample size and should be re-evaluated with larger cohorts.

The estimates of HbA1c (8) included in the meta-analysis (MCT) we used for reference were higher than our EHR-based inference. We note that we predicted HbA1c in exact periods, while the MCT method combined heterogeneous time point measurements across the different clinical trials, some listed as having up to five years of follow-up. This may suggest that the meta-analysis captured later stages in the progression of T2DM, characterized by higher HbA1c levels (30).

As demonstrated by our analysis, as well as by others (31), EHR data can support causal inference and allows replication of clinical trial results. The advantages of this approach in terms of the labor and costs required to expand evidence-based medicine are clear. As the availability of EHR data increases and the many theoretical and technical challenges associated with detecting and correcting for confounders are addressed, we expect causal inference based on observational data to become more widely used.

## Acknowledgments

AG and CY designed the experiments. AG performed the experiments. AC compiled domain expert confounders. AG, CY, AC and YG wrote the manuscript. AG and CY are guarantors of the paper. We would like to thank Omer Weissbrod for helpful inputs and suggestions and Michal Ozery-Flato for help with feature engineering tasks. The authors declare no conflict of interests.

The Corresponding Author has the right to grant on behalf of all authors and does grant on behalf of all authors, a worldwide licence to the Publishers and its licensees in perpetuity, in all forms, formats and media (whether known now or created in the future), to i) publish, reproduce, distribute, display and store the Contribution, ii) translate the Contribution into other languages, create adaptations, reprints, include within collections and create summaries, extracts and/or, abstracts of the Contribution, iii) create any other derivative work(s) based on the Contribution, iv) to exploit all subsidiary rights in the Contribution, v) the inclusion of electronic links from the Contribution to third party material where-ever it may be located; and, vi) licence any third party to do any or all of the above.

## Appendix: Supporting Tables and Figures

**Table S1.**
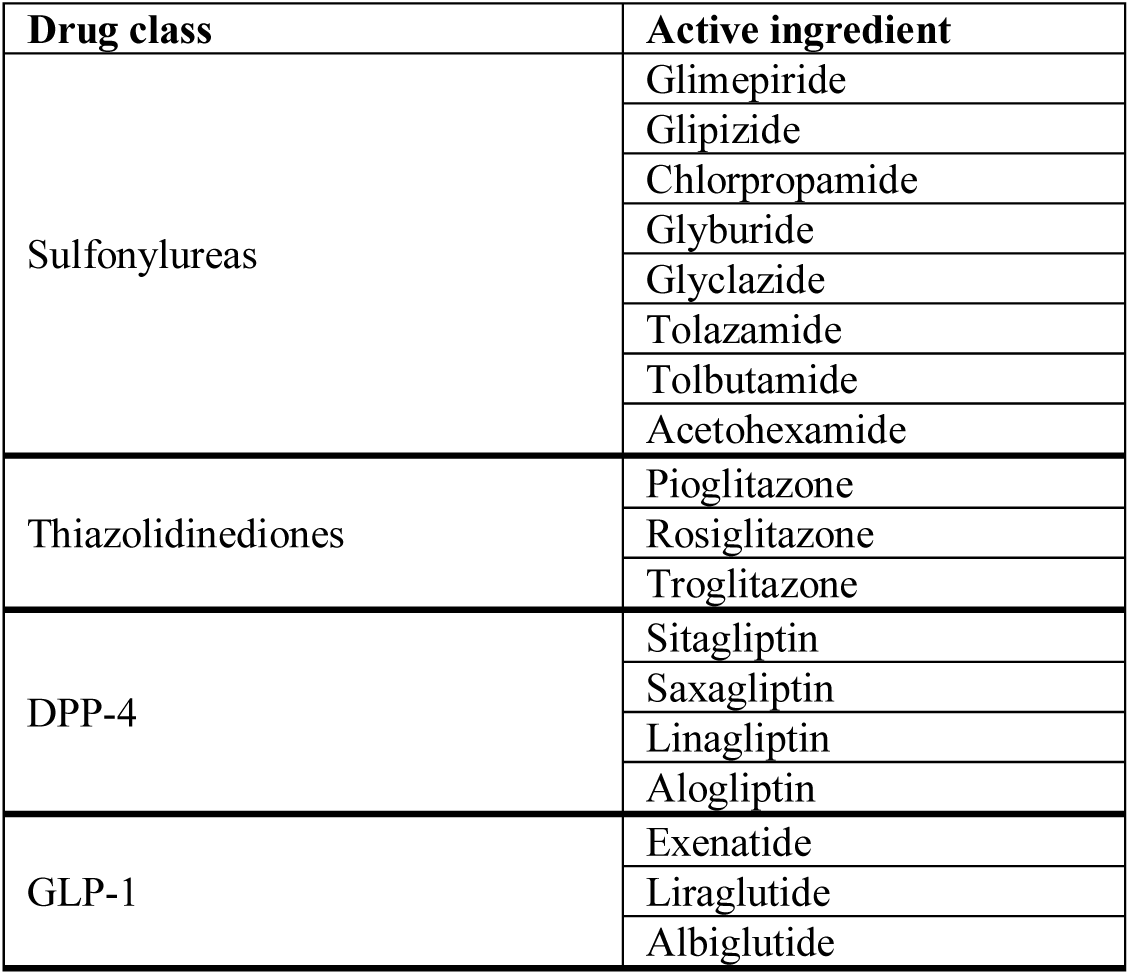
Drugs included in the compared drug classes. Missing drugs like the TZD drug lobeglitazone or GLP-1 drugs lixisenatide and dulaglutide were not prescribed to any patient in our data.

| Drug class | Active ingredient |
| --- | --- |
| Sulfonylureas | Glimepiride |
|  | Glipizide |
|  | Chlorpropamide |
|  | Glyburide |
|  | Glyclazide |
|  | Tolazamide |
|  | Tolbutamide |
|  | Acetohexamide |
| Thiazolidinediones | Pioglitazone |
|  | Rosiglitazone |
|  | Troglitazone |
| DPP-4 | Sitagliptin |
|  | Saxagliptin |
|  | Linagliptin |
|  | Alogliptin |
| GLP-1 | Exenatide |
|  | Liraglutide |
|  | Albiglutide |

**Table S2.** Selected confounders for HbA1c by domain expertise and literature (6),(13)

| Confounder category | Confounder name |
| --- | --- |
| **Demographics** | Gender |
|  | Age |
| **Vitals** | BMI |
|  | Diastolic and systolic blood pressure |
| **Prior conditions** | Dementia or Alzheimer's disease (CCS category) |
|  | Coronary or atherosclerosis condition (CCS category) |
|  | Ischemic heart disease |
|  | Diabetes neuropathy |
|  | Diabetes retinopathy |
|  | Diabetes nephropathy |
|  | Myocardial infarction |
|  | Cardiac arrest |
|  | Palliative care |
|  | History of noncompliance with medical treatment |
|  | Bed confinement |
|  | Syncope |
|  | Dizziness |
|  | Irritable colon / Irritable bowel syndrome (CCS category) |
|  | Osteoporosis |
|  | Hypertension |
|  | Hyperlipidemia |
|  | Hyperglycemia |
| **Lab values** | Baseline HbA1c measurement before second line prescription |
|  | Creatinine in blood |
|  | Total cholesterol and HDL cholesterol in blood |
|  | Triglycerides in blood |
|  | Albumin and microalbumin in urine |
|  | Protein in blood |
|  | Duration of T2DM |
| **Time to events** | Duration of metformin use |
|  | # of days between last HbA1c measurement and second line prescription |

**Table S3.** Selected confounders for BMI by a domain expert

| Confounder category | Confounder name |
| --- | --- |
| **Demographics** | Gender |
|  | Age |
| **Vitals** | BMI |
| **Prior conditions** | Concomitant SSRI, SNRI or tricyclic antidepressants |
|  | Bariatric surgery |
| **Lab values** | TSH (blood) |

**Table S4.** Pearson correlations (ρ) between actual and inferred HbA1c measurements. P-value<e^-36^ for all comparisons.

| Drug class | Standardization, comprehensive set |  | Standardization, domain expert set |  |
| --- | --- | --- | --- | --- |
|  | 6 months | 12 months | 6 months | 12 months |
| SU | 0.43 | 0.42 | 0.34 | 0.34 |
| TZD | 0.50 | 0.48 | 0.45 | 0.43 |
| GLP-1 | 0.56 | 0.52 | 0.47 | 0.46 |
| DPP-4 | 0.44 | 0.41 | 0.33 | 0.32 |

**Table S5.**
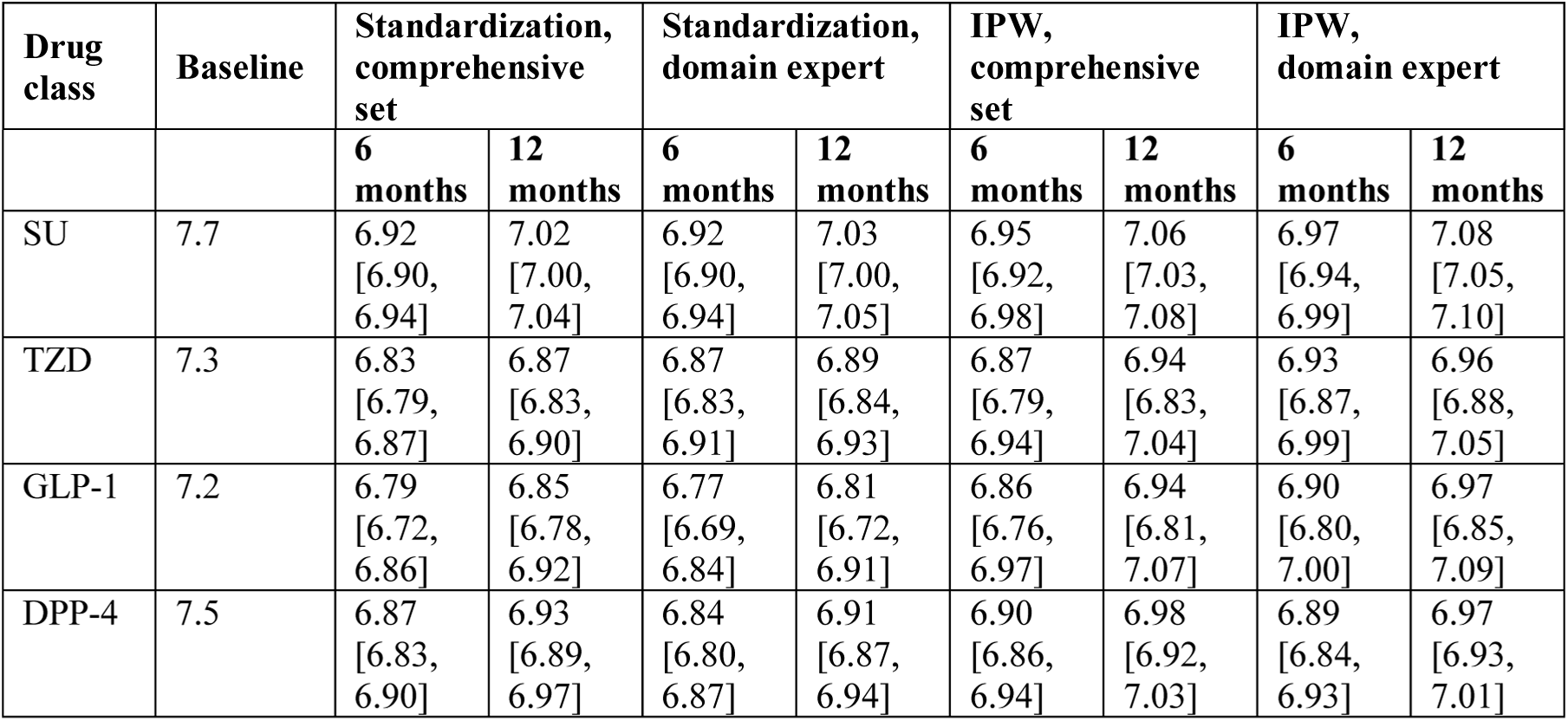
Inferred HbA1c levels in % (95% confidence intervals) for each drug class.

**Table S6.**
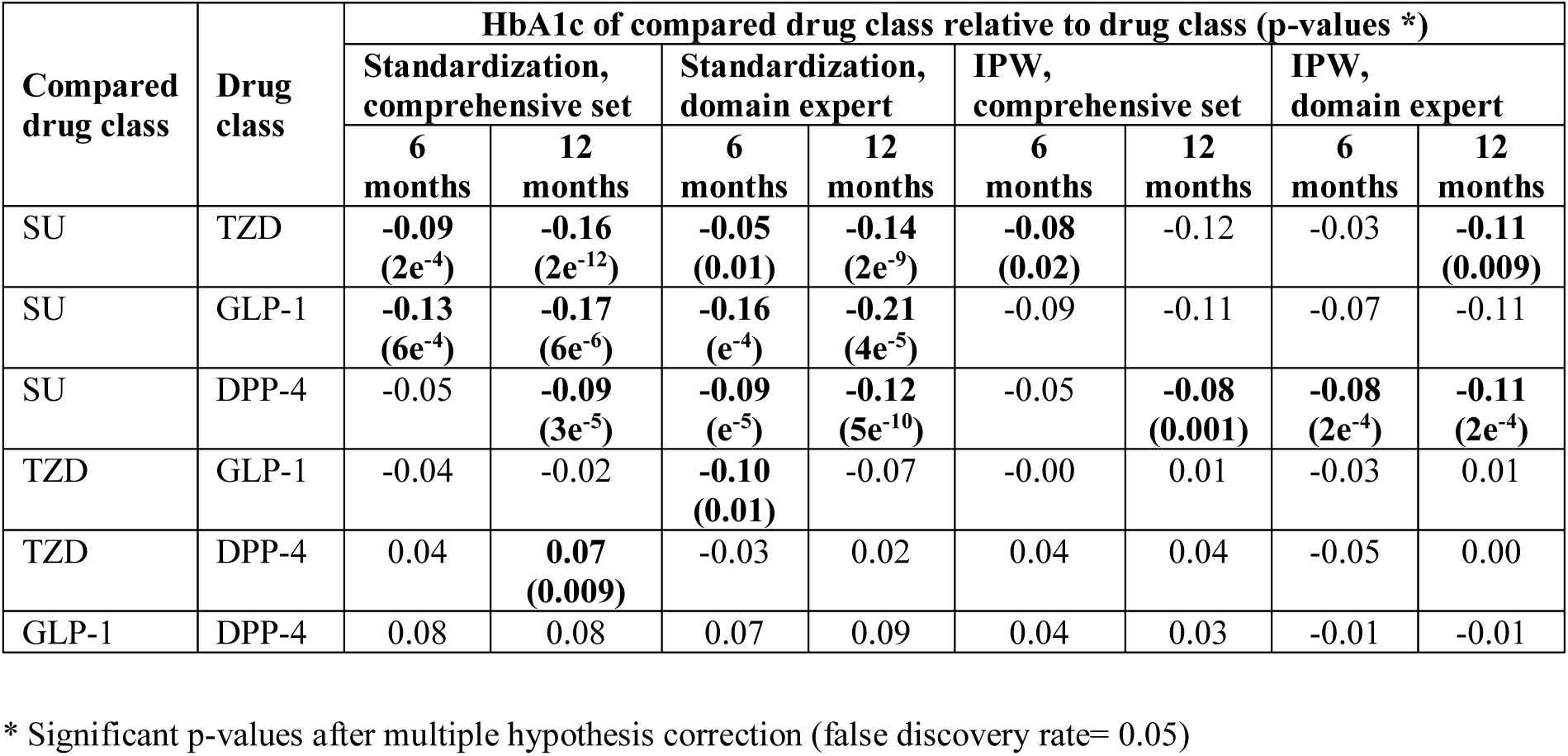
Inferred HbA1c differences between drug classes (Compared drug class serves as baseline in each row). Significant comparisons appear in bold.

**Table S7.** Inferred BMI (95% confidence intervals) for each drug class.

| Drug class | Baseline | Standardization, comprehensive set |  | Standardization, domain expert |  | IPW, comprehensive set |  | IPW, domain expert |  |
| --- | --- | --- | --- | --- | --- | --- | --- | --- | --- |
|  |  | **6 months** | **12 months** | **6 months** | **12 months** | **6 months** | **12 months** | **6 months** | **12 months** |
| SU | 33.7 | 33.8<br>[33.6, 33.9] | 33.7<br>[33.6, 33.9] | 33.8<br>[33.7, 33.9] | 33.7<br>[33.6, 33.9] | 33.9<br>[33.7, 34.2] | 33.9<br>[33.6, 34.1] | 33.8<br>[33.7, 33.9] | 33.8<br>[33.6, 33.9] |
| TZD | 33.8 | 34.0<br>[33.6, 34.3] | 34.0<br>[33.7, 34.3] | 34.2<br>[33.9, 34.4] | 34.2<br>[33.9, 34.5] | 33.5<br>[32.9, 34.1] | 33.6<br>[33.0, 34.2] | 34.0<br>[33.7, 34.3] | 34.0<br>[33.7, 34.3] |
| GLP-1 | 37.4 | 33.8<br>[33.4,<br>34.2] | 33.7<br>[33.2,<br>34.2] | 33.5<br>[33.2,<br>33.9] | 33.4<br>[33.0,<br>33.8] | 36.0<br>[35.4,<br>36.6] | 35.9<br>[35.2,<br>36.5] | 35.5<br>[35.0,<br>36.0] | 35.3<br>[34.9,<br>35.8] |
| DPP-4 | 33.9 | 33.4<br>[33.2,<br>33.6] | 33.4<br>[33.1,<br>33.6] | 33.4<br>[33.2,<br>33.5] | 33.3<br>[33.1,<br>33.4] | 33.5<br>[33.2,<br>33.8] | 33.4<br>[33.1,<br>33.7] | 33.3<br>[33.1,<br>33.5] | 33.2<br>[33.0,<br>33.4] |

**Table S8.** Inferred BMI differences between drug classes.

| Compared drug class | Drug class | BMI of compared drug class relative to drug class (*) |  |  |  |  |  |  |  |
| --- | --- | --- | --- | --- | --- | --- | --- | --- | --- |
|  |  | Standardization, comprehensive set |  | Standardization, domain expert |  | IPW, comprehensive set |  | IPW, domain expert |  |
|  |  | 6 months | 12 months | 6 months | 12 months | 6 months | 12 months | 6 months | 12 months |
| SU | TZD | 0.16 | 0.25 | **0.35**<br>(0.01) | **0.43**<br>(0.005) | -0.43 | -0.27 | 0.15 | 0.24 |
| SU | GLP-1 | 0.04 | -0.01 | -0.27 | -0.33 | **2.05**<br>(4e<sup>-10</sup>) | **1.99**<br>(4e<sup>-8</sup>) | **1.67**<br>(5e<sup>-11</sup>) | **1.58**<br>(e<sup>-11</sup>) |
| SU | DPP-4 | **-0.37**<br>(0.004) | **-0.37**<br>(0.007) | **-0.45**<br>(e<sup>-9</sup>) | **-0.49**<br>(2e<sup>-11</sup>) | **-0.45**<br>(0.02) | **-0.46**<br>(0.01) | **-0.47**<br>(3e<sup>-8</sup>) | **-0.53**<br>(6e<sup>-8</sup>) |
| TZD | GLP-1 | -0.12 | -0.26 | **-0.62**<br>(0.002) | **-0.77**<br>(8e<sup>-4</sup>) | **2.48**<br>(3e<sup>-9</sup>) | **2.26**<br>(e<sup>-6</sup>) | **1.51**<br>(4e<sup>-7</sup>) | **1.34**<br>(6e<sup>-6</sup>) |
| TZD | DPP-4 | **-0.53**<br>(0.01) | **-0.61**<br>(4e<sup>-4</sup>) | **-0.80**<br>(7e<sup>-8</sup>) | **-0.93**<br>(e<sup>-7</sup>) | -0.03 | -0.19 | **-0.63**<br>(5e<sup>-4</sup>) | **-0.77**<br>(e<sup>-5</sup>) |
| GLP-1 | DPP-4 | -0.41 | -0.35 | -0.18 | -0.16 | **-2.50**<br>(7e<sup>-14</sup>) | **-2.45**<br>(e<sup>-11</sup>) | **-2.14**<br>(9e<sup>-15</sup>) | **-2.11**<br>(3e<sup>-19</sup>) |

\* In parentheses, significant p-values after multiple hypothesis correction (false discovery rate= 0.05)

**Figure S1:**
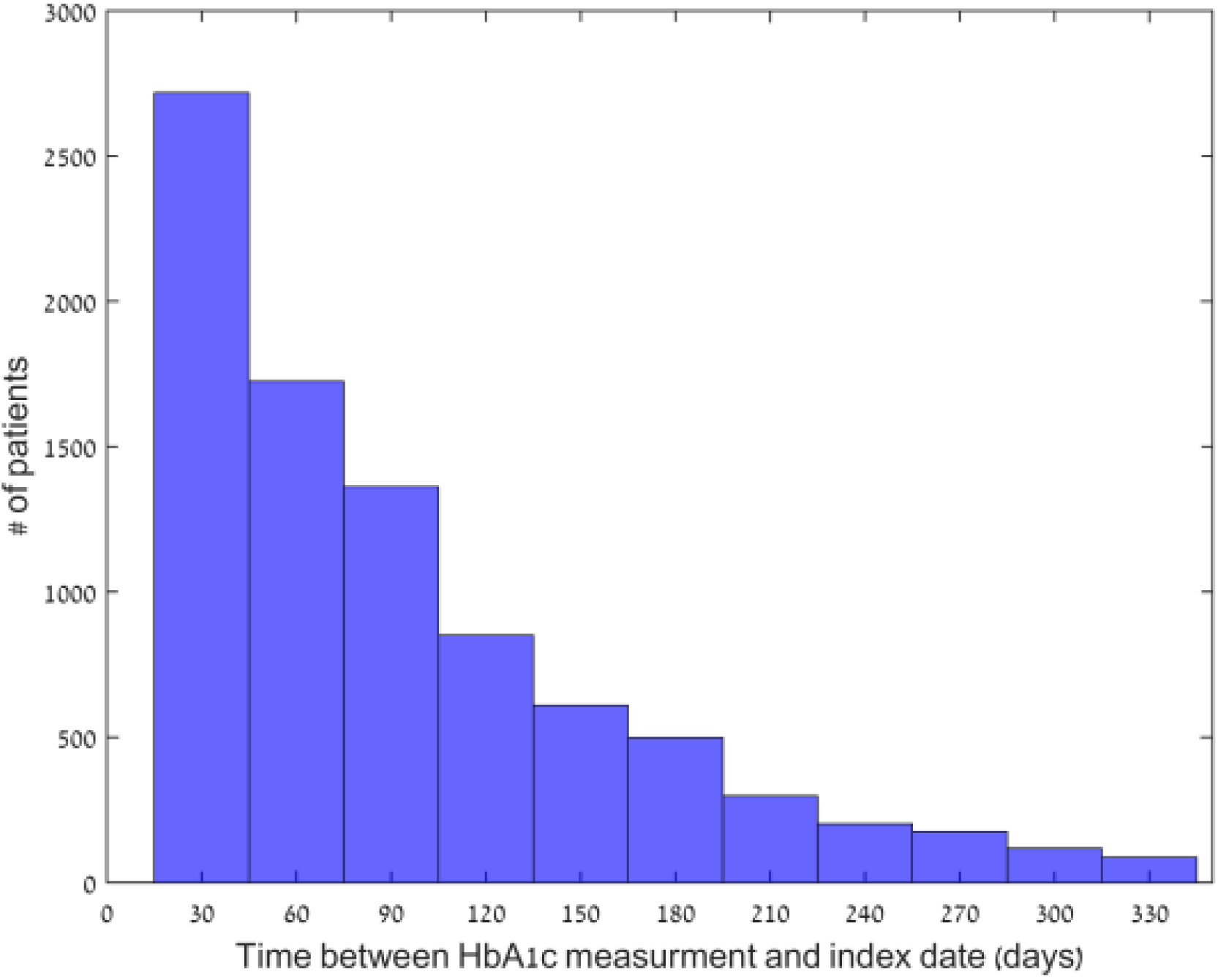
Distribution of the time difference between last baseline HbA1c measurement and initiation of second-line treatment (days).

**Figure S2:**
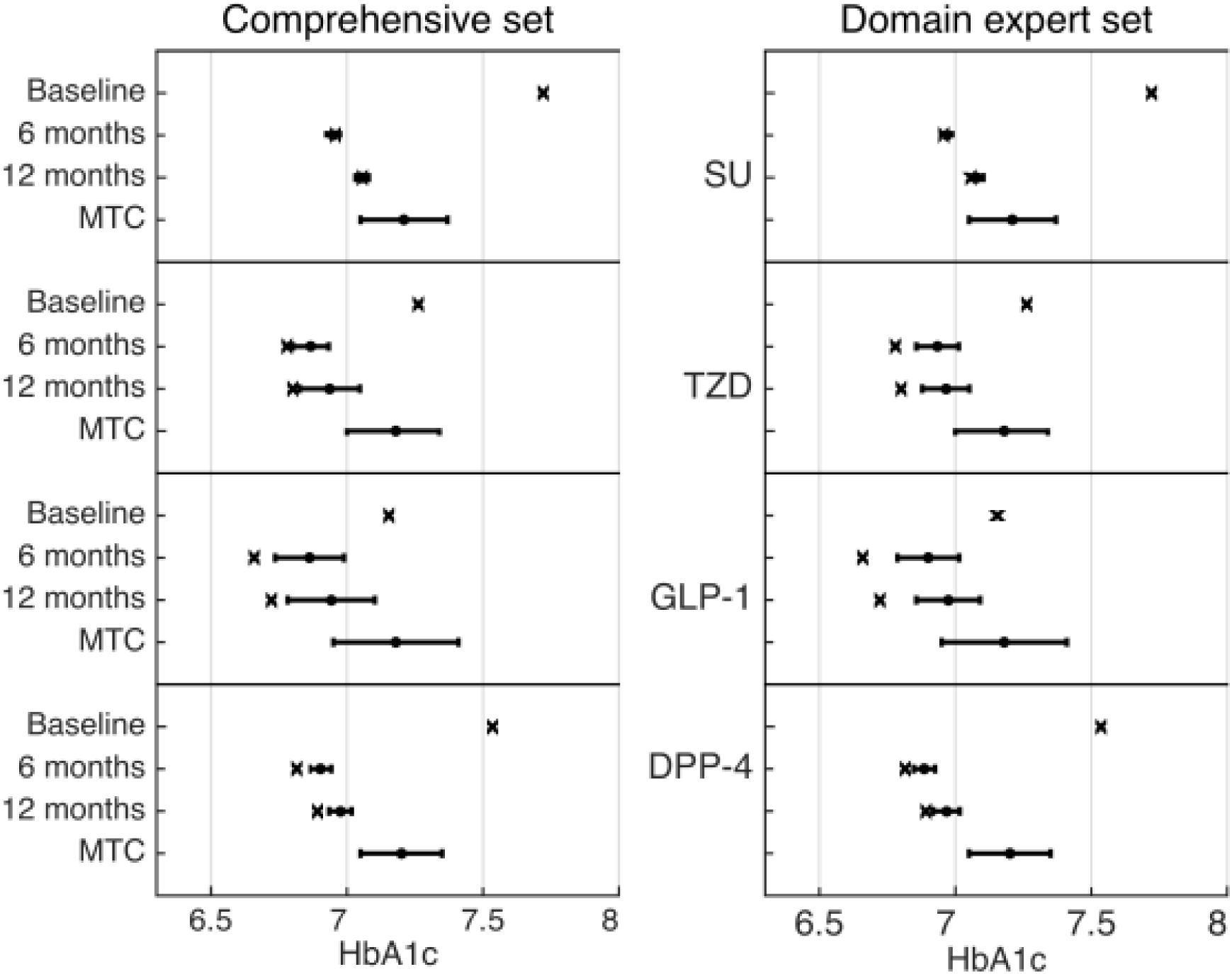
Predicted and observed HbA1c levels using inverse probability weighting adjusting for either a comprehensive set of confounders (Left panel) or a set of confounders provided by a domain expert (Right panel). X marks indicate the actual measurements of patients at baseline (before second-line treatment), after six and twelve months. Black dots (with error bars) represent the counterfactual predictions and 95% confidence intervals, supposing all patients were treated with that drug class. The results of the Bayesian mixed-treatment comparison meta-analysis by McIntosh et. al (8) are marked MTC.

**Figure S3:**
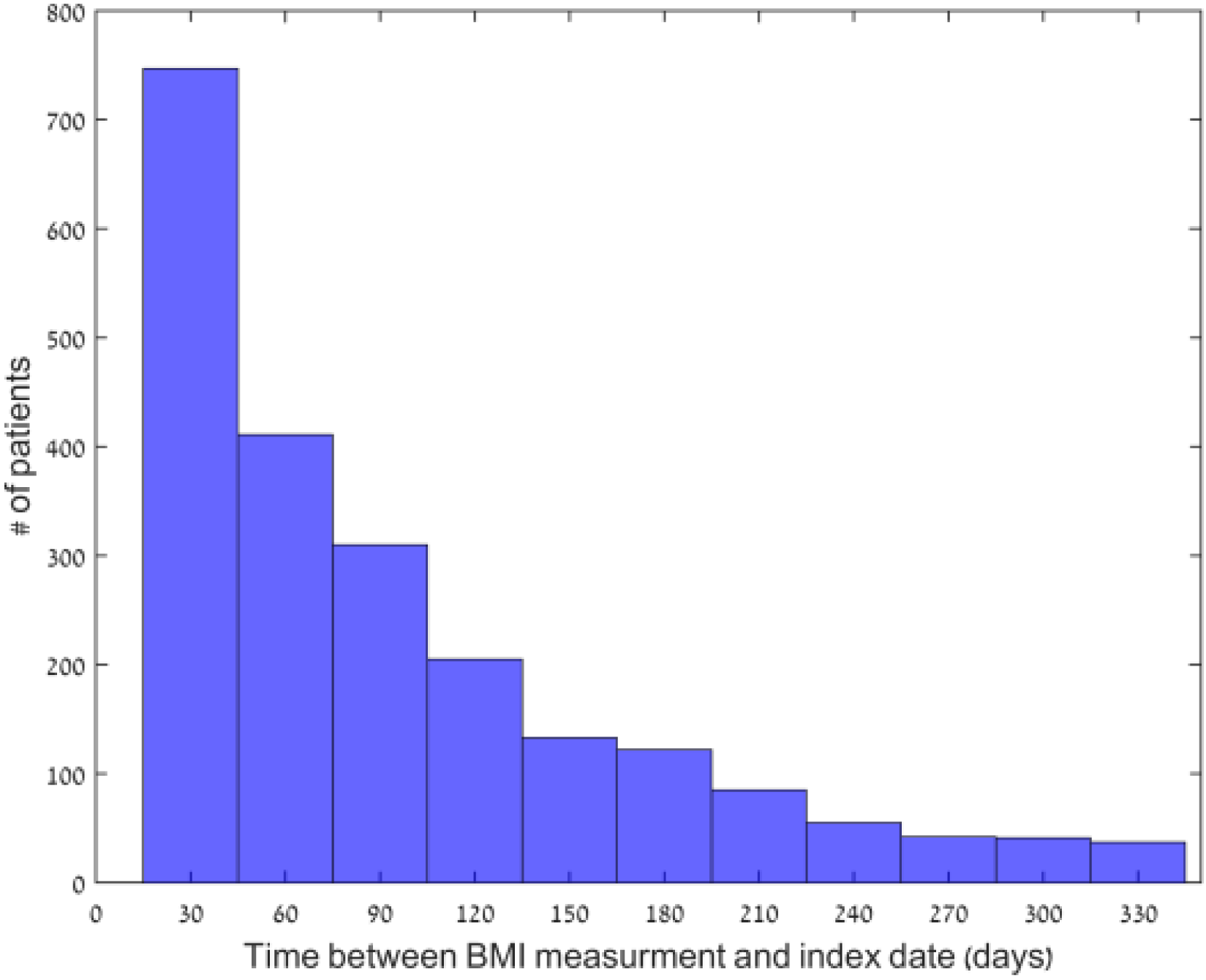
Distribution of the time difference between last BMI measurement and initiation of second-line treatment (days).

**Figure S4:**
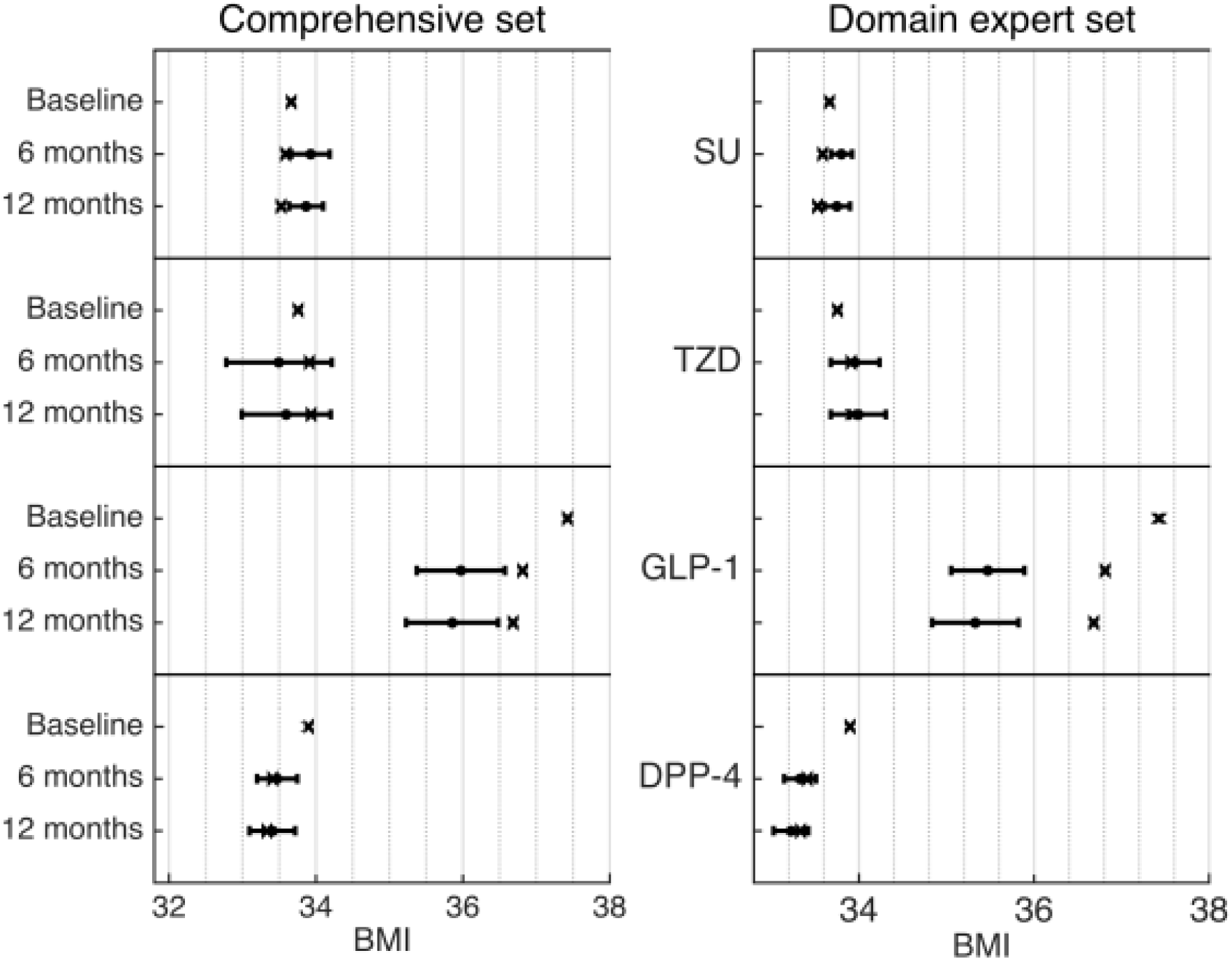
Predicted and observed BMI levels using inverse probability weighting adjusting for either a comprehensive set of confounders (Left panel) or a set of confounders provided by a domain expert (Right panel). X marks indicate the actual measurements of patients at baseline (before second-line treatment), after six and twelve months. Black dots (with error bars) represent the counterfactual predictions and 95% confidence intervals, supposing all patients were treated with that drug class

